# The gold complex auranofin sensitizes platinum resistant epithelial ovarian cancer cells to cisplatin

**DOI:** 10.1101/2024.12.03.626677

**Authors:** Farah H Abdalbari, Benjamin N Forgie, Edith Zorychta, Alicia A Goyeneche, Abu Shadat M Noman, Carlos M Telleria

## Abstract

Although there are numerous studies on drug development for ovarian cancer (OC), survival rates for this disease remain low due to platinum (Pt) resistance. Following several rounds of Pt-based chemotherapy, OC cells develop resistance by increasing DNA repair and antioxidant systems. This study aimed to design a treatment approach to combat recurrent stages of OC by repurposing the anti-rheumatic gold complex auranofin (AF). Here we demonstrate that AF enhances the efficacy of cisplatin (CDDP) in Pt-resistant epithelial OC (EOC) cells. The drug combination induces mitochondrial-dependent apoptosis, PARP cleavage, DNA damage, and ROS overproduction. These results suggest the high potential for combining AF with CDDP as a second-line therapy for recurrent EOCs.

## 1. Introduction

The survival rate for ovarian cancer (OC) patients remains at 30-40%, largely due to chemotherapy resistance [1]. The present first-line treatment for epithelial OC (EOC), cytoreductive surgery followed by platinum (Pt)-based chemotherapy, was developed in the late 1970s [2,3]. The Pt derivative cisplatin (CDDP) was approved by the FDA for the treatment of advanced EOC and bladder cancer in 1978 (revisited in [4]). CDDP induces DNA damage via the formation of crosslinks [5,6]. Other cellular targets of CDDP include the mitochondria through the induction of mitochondrial reactive oxygen species (ROS), and the plasma membrane through changes in membrane fluidity [6]. Mechanisms of resistance acquired by cancer cells against CDDP include increased DNA repair, clearance of misfolded proteins by the ubiquitin proteasome system, and reduced ROS levels via increased abundance of the antioxidant glutathione (GSH) [6]. Consequently, 80% of patients with EOC will experience disease recurrence [7]. This highlights the need for a therapeutic agent to be coupled with CDDP as a second-line therapy to increase progression-free survival for this fatal disease.

Auranofin (AF), a gold complex approved in 1985 for rheumatoid arthritis, elicits cytotoxic effects in various cancers, including some subtypes of EOC [8,9]. We previously demonstrated the potent cytotoxicity of AF against the most frequent histological subtype of EOC, high-grade serous ovarian cancer (HGSOC) [10], in which AF targets the thioredoxin reductase (TrxR) antioxidant system and induces depolarization of the mitochondrial membrane [11].

AF and CDDP target similar pathways through various mechanisms, such as overproduction of ROS and inhibition of the protein degradation system [6,12–14], suggesting that their combination may potentiate their cytotoxicity towards EOC cells.

Few studies have explored the interaction of AF and CDDP against cancer, particularly in EOC. Studies done in urothelial carcinoma [15] and small-cell lung cancer (SCLC) [16], have shown that AF increased the therapeutic effect of CDDP through cell cycle arrest and ROS accumulation [16]. Thus, we hypothesized that combining AF with CDDP in EOC cells will result in enhanced cytotoxicity compared to the individual drugs. We explored the cytotoxic efficacy of the AF/CDDP combination in two models of CDDP-resistant EOC cells.

We provide evidence that demonstrates a synergistic cytotoxic interaction between AF and CDDP in CDDP-resistant EOC cells through intrinsic caspase-dependent apoptosis, DNA damage, mitochondrial membrane depolarization, and oxidative stress. In other words, we demonstrated that the two metal-derived drugs are better than one in triggering cell death in EOC.

## 2. Materials and methods

### 2.1. Reagents and cell lines

TOV112D cells were isolated from a 42-year-old patient with grade 3 endometrioid carcinoma and are spontaneously resistant to clinically achievable concentrations of CDDP [17,18]; they were reclassified as epithelial endometrioid ovarian carcinoma based on extensive transcriptional profiling [17]. IGROV-1/CP is a cell line derived from IGROV-1 cells which had acquired resistance to CDDP by prolonged exposure in culture to the Pt drug [19]; the original IGROV-1 cells were isolated from a grade 3 solid primary ovarian tumour of a 47-year-old patient [20] with a diagnose of clear-cell ovarian carcinoma [17,21]. The two cell lines were cultured in RPMI 1640 media powder (Gibco™, Hampton, New Hampshire, USA) dissolved in HyPure™ Cell Culture Grade Water (Cytiva, Marlborough, Massachusetts, USA). The composition of the media was reported in detail recently [11]. The drugs used in this study include the following: auranofin (AF; A6733, Sigma Chemical Co., St. Louis, MO, USA), *cis-* diammineplatinum (II) dichloride (cisplatin [CDDP], P4394, Millipore, Burlington, Massachusetts, USA), and N-acetyl-L-cysteine (NAC) (A7250, Sigma-Aldrich, St. Louis, Missouri, USA).

### 2.2. Measuring IC50s and combination indexes

The concentration of CDDP that kills cells by 50% (IC50) with or without AF was determined based on cellular viability. IGROV-1/CP or TOV112D were incubated with increasing concentrations of CDDP for 3 h followed by 72-h exposure or not to 2 µM AF. The viability was measured using the Muse® count and viability reagent (MCH600103, Luminex, Austin, TX, USA) [11]. The combination index (CI), which signifies the drug interaction between CDDP and AF, was calculated using the Chou and Talalay method [22]; a CI>1 is considered antagonistic, a CI=0 indicates no drug interaction, a CI=1 is considered additive, and a CI<1 signifies synergism.

### 2.3. Annexin-V staining

IGROV-1/CP and TOV112D were treated with 10 μM CDDP for 3 h followed by a 72-h incubation in the presence or absence of 2 μM or 0.5 μM AF, respectively. Cells were stained with Annexin V and 7-AAD (Luminex). Annexin V is expressed on the extracellular surface of the plasma membrane when the cell undergoes early apoptosis, whereas 7-AAD passes the plasma membrane into the cytoplasm due to plasma membrane rupture marking late apoptosis. Details of this technique are reported in our previous publication [11].

### 2.4. Cell cycle analysis

IGROV-1/CP cells and TOV112D cells were treated with 10 µM CDDP for 3 h followed by a 72-h treatment with 2 µM or 0.5 µM AF, respectively. Cell cycle distribution was assessed as previously described by microcytometry [11,23].

### 2.5. Analysis of the mitochondrial membrane potential

IGROV-1/CP cells were treated with 15 µM CDDP for 3 h followed by a 72-h exposure to 2 µM AF, whereas the TOV112D were treated with 10 µM CDDP for 3 h followed by a 72-h exposure to 0.5 µM AF. The cells were stained with a working solution consisting of the cationic lipophilic dye, Mito-Potential (Luminex), followed by 7-AAD. This protocol was outlined in detail previously [11].

### 2.6. Isolation of the cytosol for the detection of cytochrome c

IGROV-1/CP cells were treated for 3 h with 10 µM CDDP followed by a 48-h exposure to 2 µM AF. We followed the specific protocol for the isolation and measurement of cytochrome c as developed by others [24].

### 2.7. Protein expression measurement

IGROV-1/CP and TOV112D were treated with 10 µM CDDP alone for 3 h, and 2 µM or 0.5 µM AF for 72 h, respectively, or the combination of CDDP for 3 h and AF for 72 h. The method of preparation, extraction, and quantification of the protein lysates was previously outlined in detail [11]. The primary antibodies used were monoclonal anti-beta actin clone AC-15 (A5442,

Sigma), polyclonal anti-PARP (9542S, Cell Signaling Technology, Danvers, MA, USA), monoclonal anti-phospho H2AX (Ser139) (9718, Cell Signaling), poly-clonal caspase-3 antibody (9662, Cell Signaling), monoclonal caspase-9 antibody (9508, Cell Signaling), monoclonal caspase-7 antibody (D2Q3L1) (12827S, Cell Signaling, polyclonal cytochrome c antibody (4272, Cell Signaling), and monoclonal ubiquitin antibody (58395S, Cell Signaling). Secondary antibodies used were goat anti-rabbit IgG (H + L) conjugate (1706515, BioRad, City, State, Country) and goat anti-mouse IgG (H + L)-HRP conjugate (1706516, BioRad).

### 2.8. Treatment with the caspase-inhibitor z-DEVD-fmk

IGROV-1/CP and TOV112D cells were exposed for 3 h to 10 µM CDDP and for 48 h to 2 µM AF or 0.5 µM AF, respectively, in the presence or absence of 100 µM z-DEVD-fmk (S7312, Selleck Chemicals, Houston, USA). Z-DEVD-fmk is an irreversible inhibitor of caspase-3, which also potently inhibits caspase-6, -7, -8, and -10 (reviewed in [25]). The cell viability was assessed using the Guava Muse® cell analyzer (Millipore).

### 2.9. Measurement of intracellular levels of reactive oxygen species

IGROV-1/CP cells were treated 3 h with 15 µM CDDP followed by 72-h exposure to 2 µM AF, whereas TOV112D cells were exposed to 10 µM CDDP for 3 h followed by 48-h exposure to 0.5 µM AF. Superoxide levels were detected using an oxidative stress assay previously described in detail [11].

### 2.10. Treatment with antioxidant NAC

To determine whether the cytotoxicity induced by the combination of AF and CDDP was ROS-dependent, IGROV-1/CP cells and TOV112D cells were treated with 10 µM CDDP for 3 h and 2 µM AF or 0.5 µM AF, respectively for 48 h in the presence or absence of 5 mM N-acetyl cysteine (NAC) (Sigma). Cell viability was determined using the Guava Muse® cell analyzer (Millipore).

## 3. Results

### 3.1. AF re-sensitizes different subtypes of EOC cells to CDDP

To determine whether AF increases the sensitivity of Pt-resistant EOC cells to CDDP, we exposed IGROV-1/CP cells or TOV112D cells to a fixed concentration (2 µM) of AF and increasing concentrations of CDDP. In IGROV-1/CP cells, the concentration of CDDP that killed 50% of the cells (i.e., the IC50) was of 200 ± 23.4 μM (n=3), which decreased to the clinically achievable concentration of 17 ± 3.12 µM (n=3) following the addition of AF. In TOV112D cells, the IC50 for CDDP was 80 ± 23.6 µM (n=3), which decreased to 25 ± 1.1 µM (n=3) in the presence of AF (Fig.1A). Furthermore, the interaction between the various concentrations of CDDP and the fixed concentration of AF in both cell lines was pharmacologically synergistic, with CI values below 1 (Fig.1B).

**Fig. 1.**
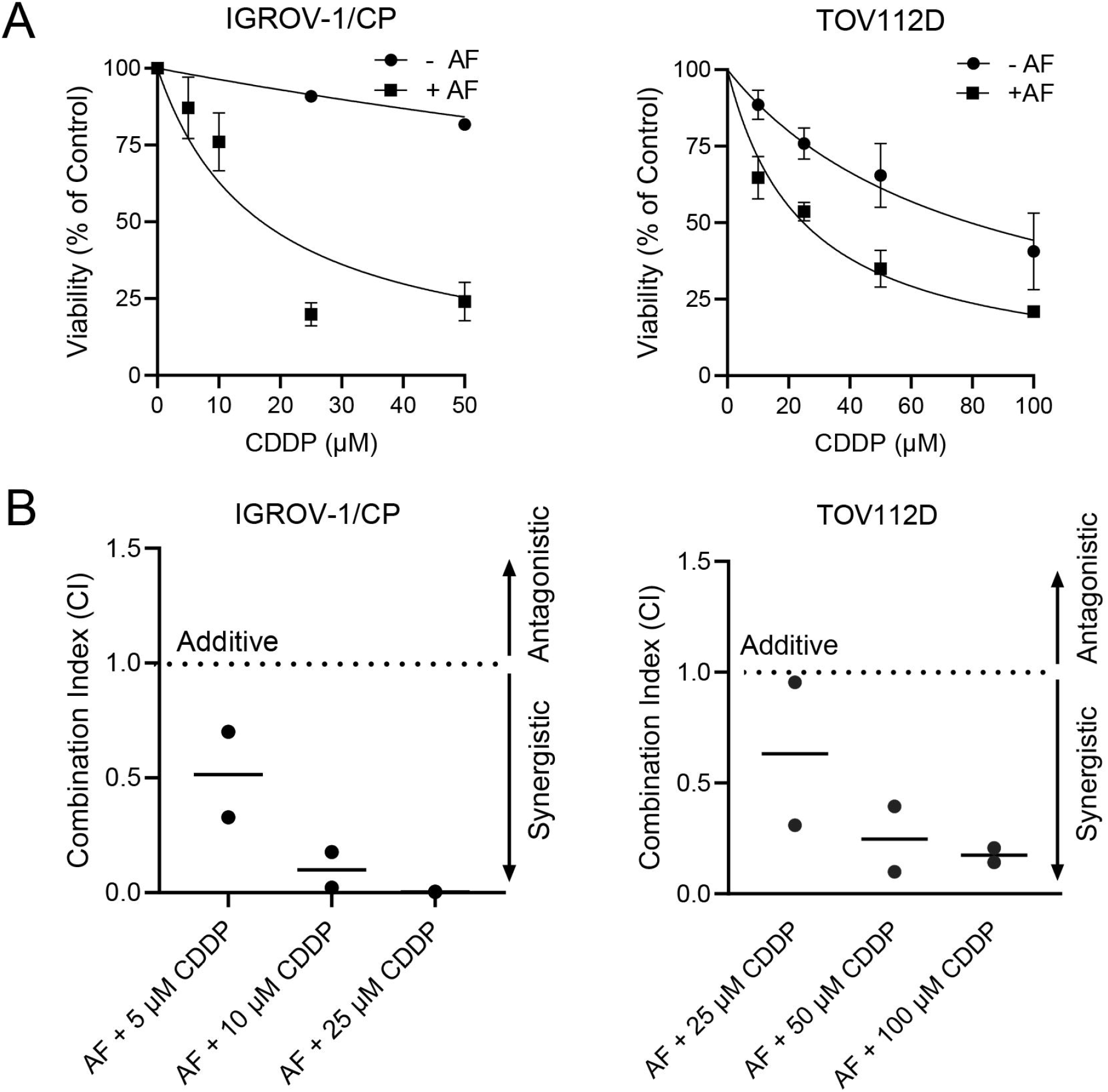
Effect of AF on the sensitization of IGROV-1/CP and TOV112D cells to CDDP. Cells were treated with increased concentrations of CDDP for 3 h followed or not by a fixed concentration (2 µM) of AF for 72 h. The viability was determined using microcytometry. Panel A shows a decrease in the IC50s of CDDP in both cell lines following the addition of AF. Panel B displays the synergistic interaction between AF and CDDP (CI<1) in the two cell lines.

### 3.2. The cytotoxicity of AF and CDDP against EOC occurs via apoptosis

The cytotoxic effect of the combination of AF and CDDP against EOC was associated with an increase in the marker of apoptotic cell death, Annexin V (Fig.2A). CDDP alone was able to induce sufficient Annexin V labelling in IGROV-1/CP cells. However, in the presence of the combination of AF plus CDDP, there was enhanced Annexin V staining in these cells. In TOV112D cells, there was a slight increase in Annexin V labelling by CDDP alone, while the combination of AF and CDDP elicited enhanced Annexin V labeling suggesting apoptotic cell death. AF alone failed to induce apoptosis in either of the cell types. In addition to the increased Annexin V labelling, the drug combination caused an accumulation of hypodiploid DNA content in both IGROV-1/CP cells and TOV112D cells (Fig.2B) further suggesting the occurrence of apoptotic cell death [26].

**Fig. 2.**
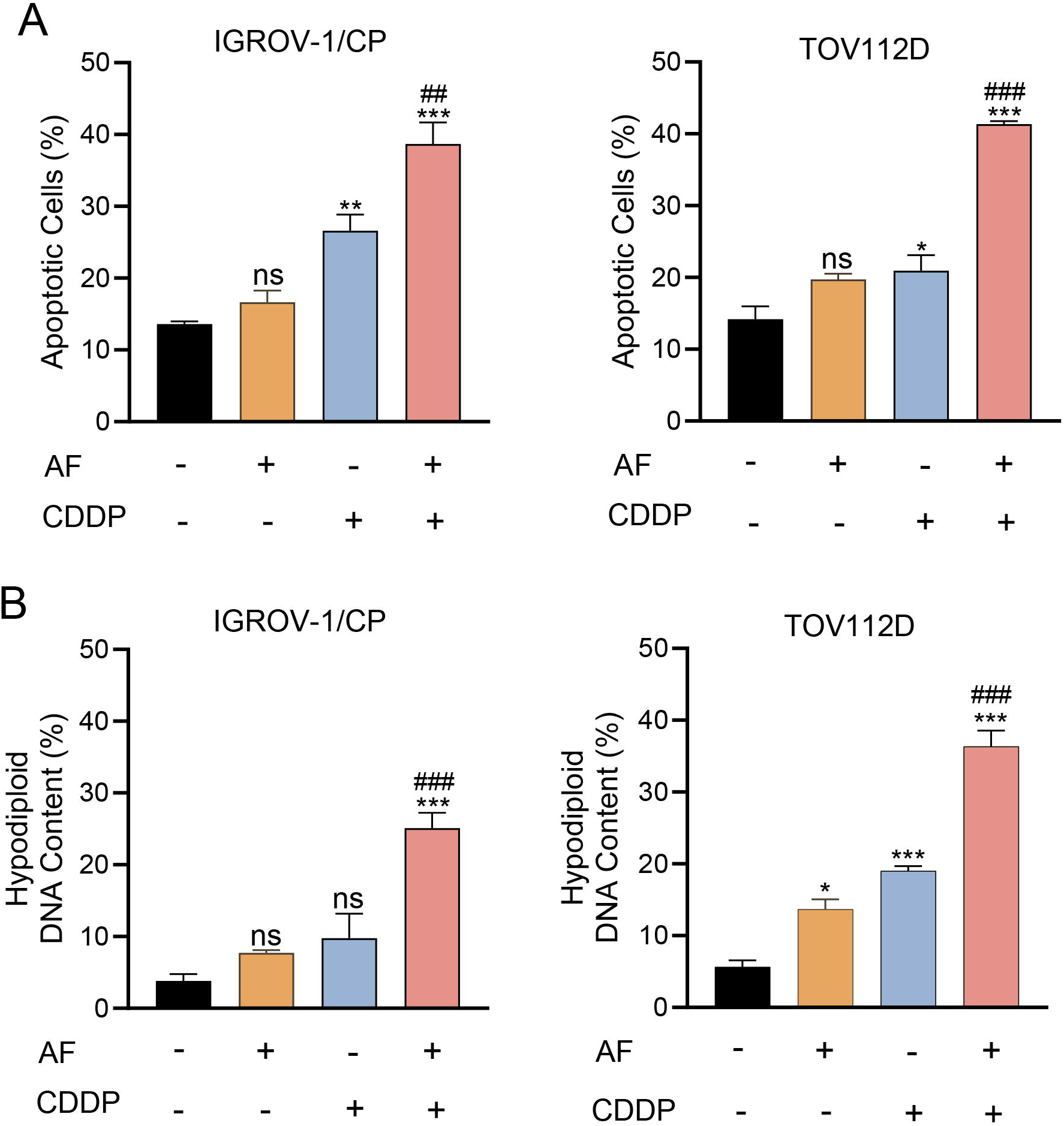
Effect of AF/CDDP combination on the induction of apoptosis in IGROV-1/CP and TOV112D cells. IGROV-1/CP and TOV112D cells were treated with 10 µM CDDP for 3 h followed by a 72-h exposure to 2 µM or 0.5 µM AF, respectively; at the end of the incubation cells were stained either with Annexin V and 7-AAD to determine apoptosis (Panel A) or propidium iodide to assess cell cycle analysis (Panel B). * p < 0.05, ** p < 0.01 and *** p < 0.001 when compared to vehicle. ## p < 0.01 and ### p < 0.001 when compared to AF- or CDDP-treated cells.

### 3.3. The cytotoxicity caused by the combination AF/CDDP is associated with intrinsic apoptosis, PARP cleavage, and DNA damage

To further confirm that apoptosis induced by the combination AF/CDDP in IGROV-1/CP and TOV112D cells is mediated by the intrinsic or mitochondrial pathway, we assessed the cleavage of the initiator caspase-9. Caspase-9 activation by the combination AF/CDDP was evident by the cleavage of the pro-caspase form into fragments weighing 49, 39, and 37 kilodaltons (kDa). Interestingly, the cleavage of the pro-caspase-9 form was also induced by CDDP alone in both cell types (Fig.3A).

**Fig. 3.**
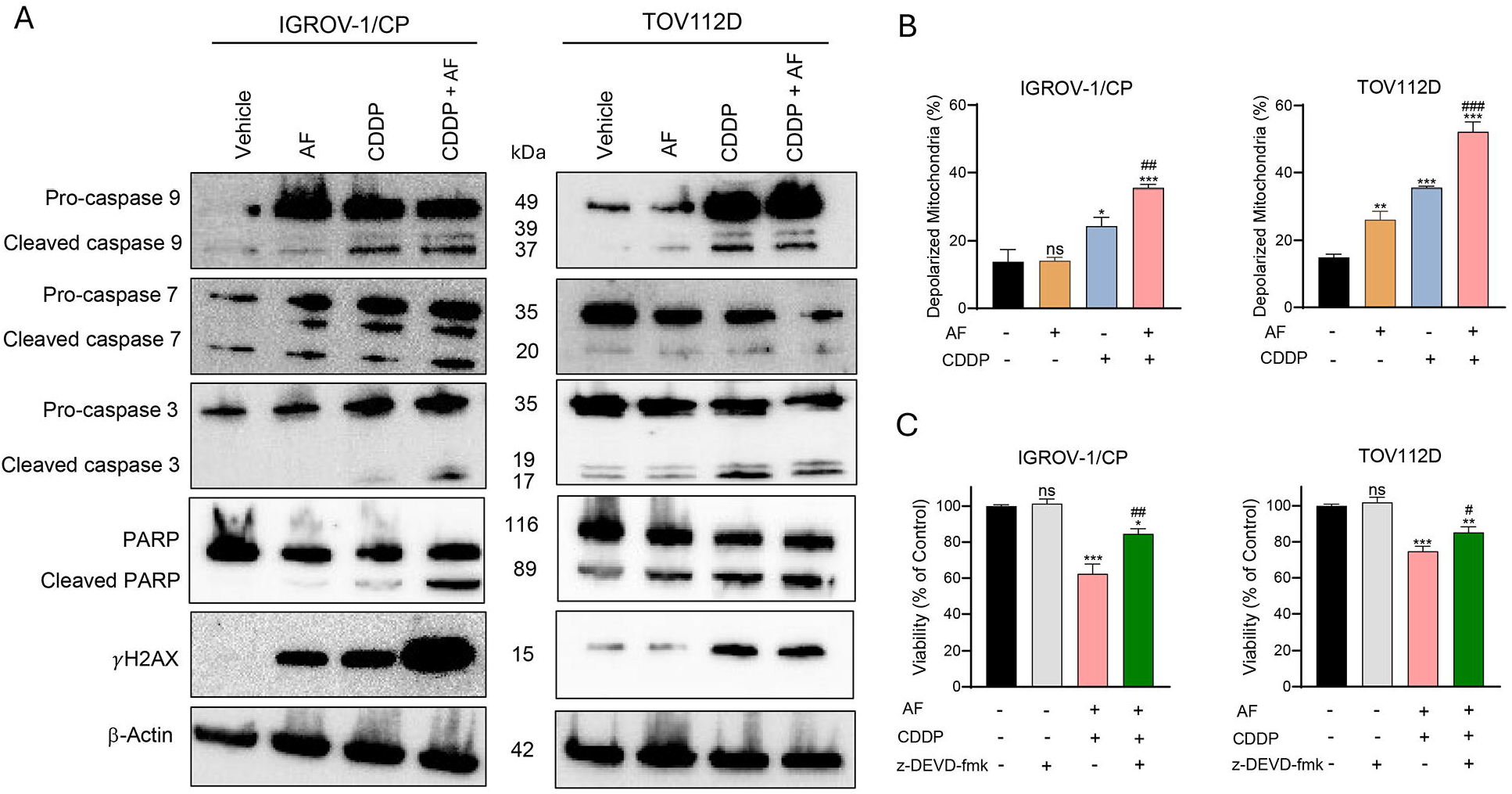
A) Effect of CDDP (3 h), AF (72 h), or their combination on the cleavage of caspases-9, - 7, and -3, and PARP, and the expression of *γ*H2AX in IGROV-1/CP and TOV112D cells as detected by western blotting. *β*-actin was used as a protein loading control. B) AF/CDDP combination increases mitochondrial membrane depolarization in comparison to each drug alone. * p < 0.05, ** p < 0.01, and *** p < 0.001 compared to vehicle; ### p < 0.001 when compared to each drug separately. C) The caspase-3 inhibitor z-DEVD-fmk partially rescues the cells from AF/CDDP-induced cytotoxicity following 48 h treatment; * p <0.05, ** p <0.01, *** p < 0.001 when compared to vehicle. # p < 0.05 and ## p < 0.01 when compared to AF/CDDP treatment.

Caspases-3 and -7 are executioner caspases in the apoptotic pathway, which target or cleave similar substrates during the programmed cell death pathway [27]. In the IGROV-1/CP cells, slight cleavage of caspases-3 and -7 was induced by CDDP alone, whereas the cleavage of both caspases was enhanced by the combination AF/CDDP. In agreement with the induction of apoptosis by the drug combination, we measured the cleavage of poly ADP-ribose polymerase-1 (PARP), a nuclear enzyme that is heavily involved in DNA repair [28]. PARP can be cleaved into 89 kDa and 24 kDa fragments by caspases-3 and -7, which causes PARP to be no longer active in the DNA repair pathway [28]. In IGROV-1/CP, PARP cleavage was highly evident only by the combination of AF and CDDP. Interestingly, PARP cleavage is apparent in TOV112D cells following treatment with either AF or CDDP alone; however, it is enhanced by the combination AF/CDDP (Fig.3A).

Early events of the DNA damage response, including the cleavage of PARP, also involve the phosphorylation of histone H2AX at serine-139, resulting in the formation of *γ*H2AX. The expression of *γ*H2AX is thought to increase DNA accessibility by DNA repair enzymes [29]. As anticipated by PARP cleavage, increased DNA damage was induced by the AF/CDDP combination in IGROV-1/CP and TOV112D cells (Fig. 3A).

Since we observed the induction of mitochondrial-dependent apoptosis via caspase-9 cleavage by the AF/CDDP combination, we decided to explore the state of the mitochondria in response to the treatment. We found clear changes in the mitochondrial membrane potential of IGROV-1/CP and TOV112D cells following exposure to AF/CDDP when compared to each drug alone (Fig.3B). The mitochondrial membrane was depolarized slightly by CDDP and more dramatically by the addition of AF to CDDP in IGROV-1/CP cells. In TOV112D cells, there was a slight depolarization of the mitochondrial membrane caused by AF, more so by CDDP, and even more enhanced by the combination AF/CDDP.

Finally, we observed that the cytotoxicity induced by AF and CDDP in IGROV-1/CP cells and TOV112D cells was significantly, yet partially reversed in the presence of z-DEVD-fmk, an irreversible inhibitor of the primary executioner caspase, caspase 3 [27] (Fig. 3C).

### 3.4. AF and CDDP increase oxidative stress in EOC cells

We hypothesized that the disruption of the mitochondrial membrane potential elicited by the AF/CDDP combination in IGROV-1/CP and TOV112D cells was likely caused by ROS accumulation. To determine whether these two metals together act as pro-oxidants we measured in both cell lines the total intracellular levels of ROS in response to AF, CDDP, or the combination of AF/CDDP. In the presence of the drug combination, there was higher ROS accumulation within the TOV112D cells than that caused by each drug alone (Fig.4A). Interestingly, in IGROV-1/CP cells, there was a slight induction of ROS accumulation by AF, a further induction by CDDP, which in this case did not increase further when combined with CDDP.

**Fig. 4.**
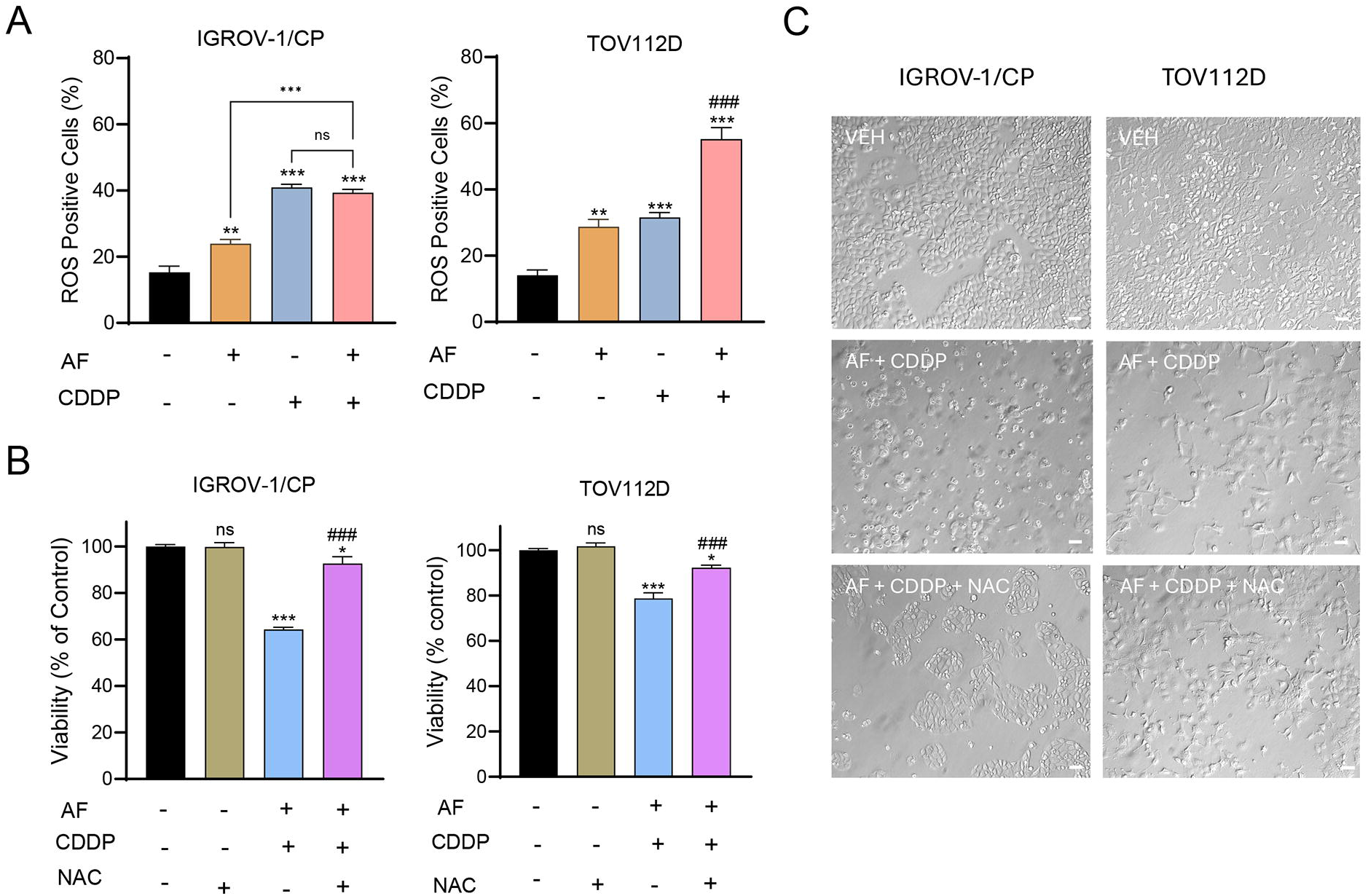
A) Effect of AF, CDDP, or their combination on the production of ROS. Treatment was performed with 15 µM CDDP for 3 h and 2 µM AF for 72 h in IGROV-1/CP cells, and 10 µM CDDP for 48 with 0.5 µM AF for 48 h in TOV112D cells. ** p < 0.01 and *** p <0.001 when compared to vehicle. ### p < 0.001 when compared to AF or CDDP alone. B) Protection of AF/CDDP-induced cytotoxicity in the presence of 5 mM NAC for 48 h. * p < 0.05 and *** p < 0.001 compared to vehicle. ### p < 0.001 when compared to AF/CDDP. C) Phase-contrast images obtained after 48 h incubation with the indicated drugs. VEH, vehicle; AF, auranofin; CDDP, cisplatin; NAC, N-acetyl-L-cysteine. Scale bars = 50 µm.

To determine whether the cytotoxic effect of the combination of AF/CDDP is dependent on ROS production, we treated the EOC cells with the drugs in the presence of the antioxidant NAC. NAC rescued IGROV-1/CP and TOV112D cells from undergoing death elicited by the drug combination (Fig. 4B). The healthy appearance of the IGROV-1/CP cells and TOV112D cells in untreated conditions changed to substantial toxicity in the presence of AF/CDDP; however, NAC was able to reverse in part the cell deterioration induced by AF/CDDP (Fig.4C).

## 4. Discussion

The present study focused on the interaction between the gold complex AF and the DNA damaging agent CDDP in Pt-resistant EOC cells. We demonstrate the existence of a synergistic drug interaction in the killing of two EOC cell types [17,19]. Despite the large differences in sensitivities to CDDP displayed by the two cell lines, the IC50s for CDDP were significantly decreased in both cell lines following the addition of AF to the range of ∼20 µM, which is within the clinically reachable concentrations [30]. A similar drug interaction was recorded in small-cell lung carcinoma cells (SCLC) [16] and urothelial carcinoma cells [31]. These findings signify that AF is a Pt sensitizer in Pt-resistant EOCs; our study is the first to explore the success of this combination therapy in this fatal disease.

Treatment with the combination AF/CDDP strongly induced apoptosis in both cell types tested. In the EOC cells we studied, apoptosis was triggered through the intrinsic pathway, as the addition of AF to CDDP enhanced caspase-9 cleavage, in addition to the executioner caspases, caspases-3 and -7 [27]. The cytotoxicity of the combination therapy in both cell types is partially dependent on caspase-3 activation, a finding that has not been reported in previous studies on this drug combination [15,16].

The combination AF/CDDP also enhanced PARP cleavage, particularly in the more Pt-resistant IGROV1/CP cells, similar to results reported in SCLC cells [16]. Additionally, DNA damage was increased in both cell lines as marked by the upregulation of γH2AX. In IGROV- 1/CP cells but not TOV112D, the addition of AF to CDDP further increased γH2AX expression.

The AF/CDDP combination treatment increased mitochondrial membrane depolarization in both cell lines, with a more pronounced effect in the less Pt-resistant TOV112D cells. This is consistent with a study showing less mitochondrial membrane depolarization in Pt-resistant OVCAR-8 cells in response to CDDP compared to CDDP-sensitive OVCAR-3 and OVCAR-4 cells [32]. Despite varying Pt sensitivities, the combination AF/CDDP successfully induced mitochondrial membrane depolarization in both cell lines studied. This depolarization likely triggered the release of cytochrome c [33], which we demonstrated in IGROV-1/CP cells subjected to cell fractionation (Supplementary Fig. 1).

Disruption of the mitochondrial membrane potential can be caused by mitochondrial ROS accumulation [6,34,35]. We previously reported that AF acts as a pro-oxidant by inhibiting the TrxR antioxidant system in HGSOC cells [11], an effect also observed in other cancers [36–38]. Similarly, CDDP generates ROS through mitochondrial damage and electron transport chain impairment [6] in lung and prostate cancer cells [39]. The AF/CDDP combination treatment enhances ROS production in TOV112D cells, but in IGROV-1/CP cells, ROS production seems to be primarily driven by CDDP. This is possibly due to upregulated antioxidant defences as a compensatory mechanism for survival [40]. Of interest, the cytotoxicity elicited by the AF/CDDP combination is ROS-dependent, as demonstrated by the rescue of viability observed when using the ROS scavenger, NAC (Fig.4B). NAC is an antioxidant via GSH replenishment or direct scavenging of oxidant species [41]. AF and CDDP have each been reported to disrupt protein homeostasis [6,9]. We observed an accumulation of poly-ubiquitinated proteins in IGROV-1/CP cells treated by the AF/CDDP, suggesting proteasome inhibition by the drug combination (Supplementary Fig.2).

In conclusion, the synergistic cytotoxic interaction of AF/CDDP in Pt-resistant EOC cells is associated with intrinsic caspase-dependent apoptosis, DNA damage, and ROS accumulation. The results imply that AF overcomes Pt resistance in EOC cells suggesting their potential utility in treating recurrent disease which almost always develops as a Pt resistant.

## CRediT authorship contribution statement

**Farah H. Abdalbari:** Investigation, Formal analysis, Methodology, Writing first draft. **Benjamin N. Forgie:** Investigation, Review and Editing. **Edith Zorychta**: Review and Editing. **Alicia A Goyeneche**: Supervision, Methodology, Review and Editing. **Abu Shadat M Noman**: Funding acquisition, Review and Editing. **Carlos M Telleria**: Conceptualization, Supervision, Resources, Project administration, Funding acquisition.

## Data availability

Data will be made available upon reasonable request.

## Acknowledgements

This work was supported by a grant from LabQuest Diagnostic Limited, Dhaka, Bangladesh and funds from the Department of Pathology, McGill University.

## Appendix A

**Supplementary data.** Supplementary data to this article can be found online at https://

## Abbreviations

EOC: Epithelial ovarian cancer
HGSOC: High-grade serous ovarian cancer
OC: Ovarian cancer
TrxR: Thioredoxin reductase
AF: Auranofin
CDDP: Cisplatin
CI: Combination index
ROS: Reactive oxygen species
NAC: N-acetyl-L-cysteine

**Pt** Platinum

