## Supplementary Figures 1 and 2 for "The gold complex auranofin sensitizes platinum resistant epithelial ovarian cancer cells to cisplatin"

**Supplementary Material**

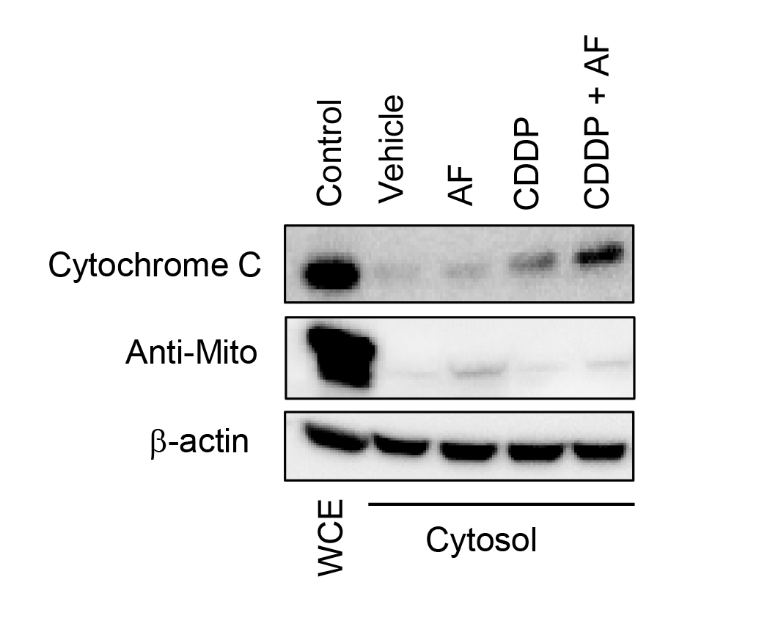

**Supplementary Fig. 1**. AF/CDDP in combination induces cytochrome c release into the cytosol, as detected by western blotting. We used whole cell extracts (WCE) to control for the presence of mitochondrial proteins in the WCE and the absence of mitochondrial proteins in the cytosol using an anti-mitochondrial antibody (Anti-Mito). It is evident how the combination of AF/CDDP enhances the release of cytochrome c into the cytosolic fraction. IGROV-1/CP cells were treated for 3 h with 10 µM CDDP, for 72 h with 2 µM AF, or with the combination of both before isolation of cytosols.

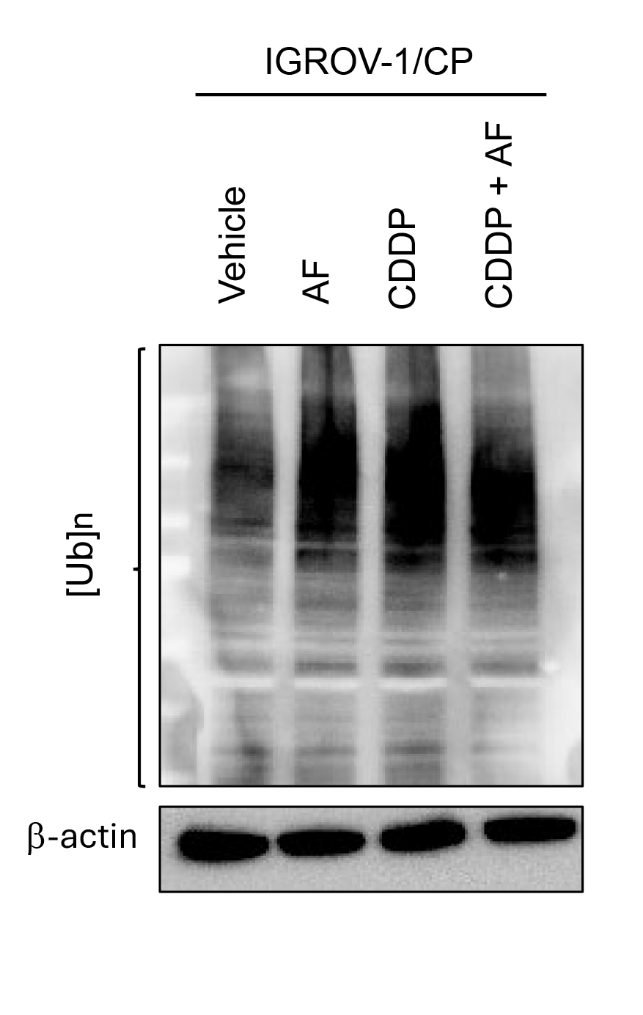

**Supplementary Fig. 2.** Accumulation of poly-ubiquitinated proteins in IGROV-1/CP cells following treatment with 10 µM CDDP for 3 h alone, with 2 µM AF for 72 h alone, or with the combination of both. [Ub]n denotes poly-ubiquitinated proteins as detected by western blotting.
